# Structure of APP-C99_1-99_ and Implications for Role of Extra-Membrane Domains in Function and Oligomerization

**DOI:** 10.1101/257469

**Authors:** George A. Pantelopulos, John E. Straub, D. Thirumalai, Yuji Sugita

## Abstract

The 99 amino acid C-terminal fragment of Amyloid Precursor Protein APP-C99 (C99) is cleaved by γ-secretase to form Aβ peptide, which plays a critical role in the etiology of Alzheimer’s Disease (AD). The structure of C99 consists of a single transmembrane domain flanked by intra and intercellular domains. While the structure of the transmembrane domain has been well characterized, little is known about the structure of the flanking domains and their role in C99 processing by γ-secretase. To gain insight into the structure of full-length C99, REMD simulations were performed for monomeric C99 in model membranes of varying thickness. We find equilibrium ensembles of C99 from simulation agree with experimentally-inferred residue insertion depths and protein backbone chemical shifts. In thin membranes, the transmembrane domain structure is correlated with extra-membrane structural states. Mean and variance of the transmembrane and G_37_G_38_ hinge angles are found to increase with thinning membrane. The N-terminus of C99 forms β-strands that may seed aggregation of Aβ on the membrane surface, promoting amyloid formation. The N-terminus, which forms α-helices that interact with the nicastrin domain of γ-secretase. The C-terminus of C99 becomes more α-helical as the membrane thickens, forming structures that may be suitable for binding by cytoplasmic proteins, while C-terminal residues essential to cytotoxic function become α-helical as the membrane thins. The heterogeneous but discrete extra-membrane domain states analyzed here open the path to new investigations of the role of C99 structure and membrane in amyloidogenesis.

## Introduction

Amyloid Precursor Protein (APP), a 770-residue membrane protein, plays important roles in neural activity and regulation of synaptic formation.[1] The canonical APP processing pathway is defined by APP cleavage by either by α- or β- secretase resulting in 83-or 99-residue long peptides (C83 and C99), that form the majority of APP fragments in cells.[2] C99 can subsequently undergo processive cleavage of its transmembrane domain by γ-secretase at various sites within its transmembrane (TM) region, yielding 38, 40, 42, and 43-residue long N-terminal fragments commonly known as Amyloid β (Aβ) protein.[3] Aβ_42_ (and to some extent Aβ_43_) has been implicated in the onset of Alzheimer’s disease (AD) due to the presence of fibrillar aggregates enriched in these peptides[4,5] found in the brains of AD patients.[6] In addition, Aβ_42_ oligomers have been directly observed to accompany loss of neural plasticity and memory.[7]

Solution NMR measurements[7,8] employing zwitterionic bicelles and micelles provide the primary source of information on the structure of C99 in a variety of membrane mimicking environments. In these *in vitro* environments, there is evidence that residues 1-14 (see Figure 1) of the N-terminal domain (NTD) are disordered, residues 15-25 of the N-terminus have helical propensity (N-Helix), residues 26-28 form a turn (N-Turn), residues 29-52 form the helical transmembrane domain (TMD), residues 53-90 of the C-terminus form a disordered region (C-Loop), and residues 91-99 of the C-terminus form a helix (C-Helix).[8,9] Insertion of residues in the membrane evidenced by EPR[8] and NMR[9] measurements suggest that in some systems the C-Helix and N-Helix domains rest on the membrane surface, while the proximities of the NTD and C-Loop domain to the membrane remain unclear.

**Figure 1.**
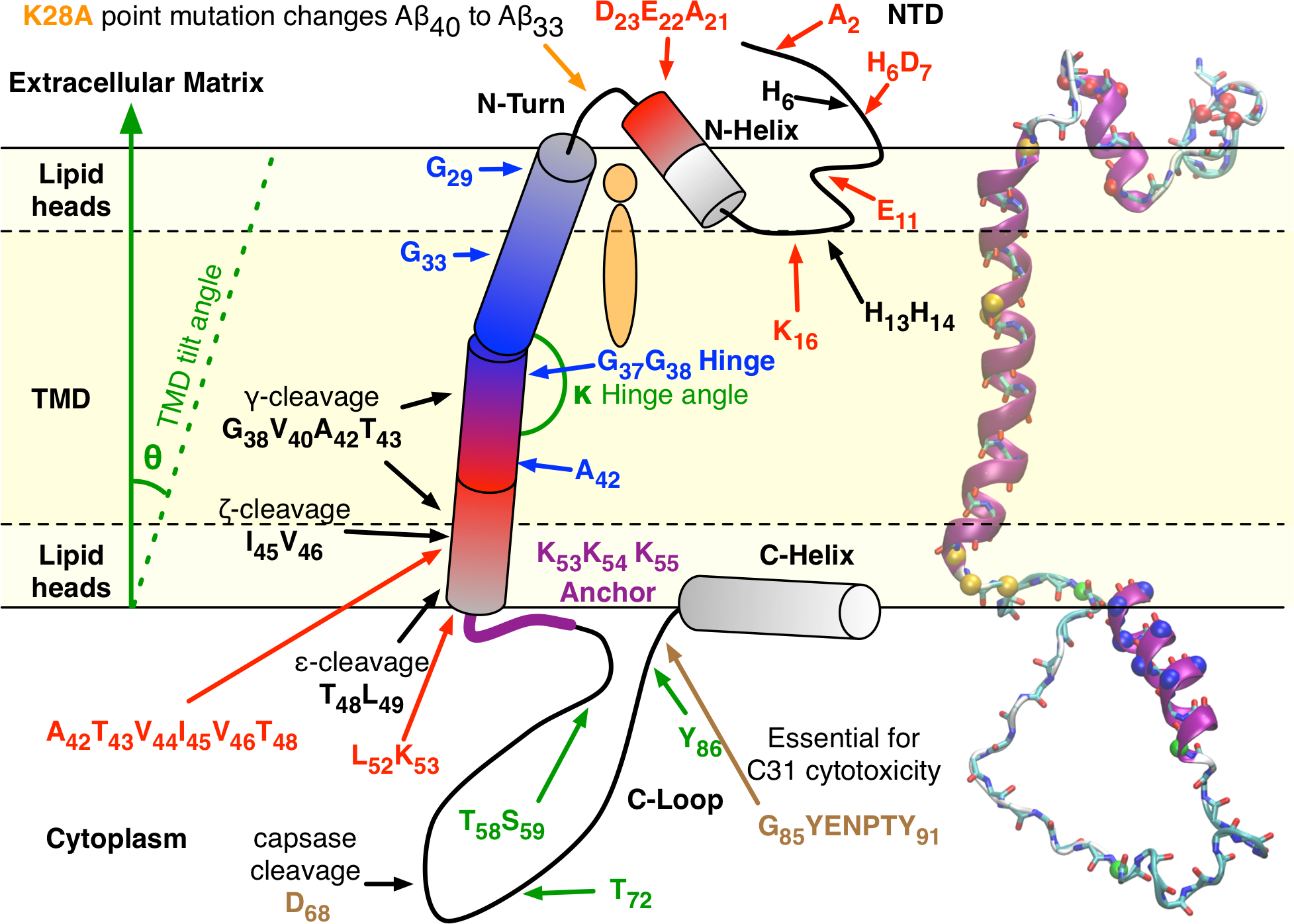
Yellow shading represents hydrophobic core of the membrane. Red residues contain familial AD mutations. Blue residues are critical to the formation of C99 dimers. Green residues may be phosphorylated. Purple residues form the lysine anchor. Black residues indicate γ-secretase cleavage sites and metal binding residues. Brown residues are critical for C31 formation and cytotoxicity. Cylinders represent domains with significant helical propensity. *θ* and *κ* angles describe the TMD tilt and GG hinge angle. The *θ* angle increases with thinning or curving of the membrane surface, and *κ* increases with curving of the membrane surface. Black solid lines mark the membrane surface and black dashed lines represent the membrane hydrophobic core. The orange lipid marks the putative cholesterol binding site. Also shown is an atomistic structure of C99 predicted from TALOS+ using LMPG micelle backbone chemical shifts and secondary structure assigned with STRIDE. Cα within the atomistic structure are labeled as N-terminal familial AD mutation (red), residues 28, 37, 38, 53,54, and 55 (orange), phosphorylatable residues Ca (green), and C-Helix (blue).

The structure of the TMD is believed to be critical to the mechanism of recognition and cleavage of C99 by γ-secretase. The process of cleavage of C99 by γ-secretase begins with the “ε-cleavage” step, forming Aβ_48_ or Aβ_49_, which are then further cleaved via “ζ-cleavage” to form Aβ_45_ and Aβ_46_. These fragments are subsequently processed by “γ-cleavage” to predominantly form Aβ_38_ or Aβ_42_, and Aβ_40_ or Aβ_43_, respectively.[10] C99 features a glycine zipper motif, G_29_xxxG_33_xxxG_37_, in the TMD, which is frequently observed in dimer-prone single-pass TM proteins.[11,12] It is further evidenced to be a component of putative cholesterol binding site on C99,[13,14] a finding that is important because cholesterol has been hypothesized to recruit C99 to γ-secretase.[14–16] Mutation of G_29_ and G_33_ in this motif reduces Aβ_42_ production,[17] and is expected to reduce C99 dimerization.[18] Proximate to the N-terminal portion of the GxxxG repeat motif lies a “GG hinge” at G37G38 in the TMD, previously identified by molecular dynamics simulations[19,20] and conjectured to be important to processing by γ-secretase.[21] Hydrogen-deuterium (H-D) exchange studies observed side chain[21] and a-helix[22] hydrogen bonds to be substantially weaker near the GG hinge, suggesting the amide bonds are readily available for γ-cleavage. Thickening of the membrane reduces the relative amount of Aβ_42_ and Aβ_43_ produced while leading to an overall increase in γ-secretase activity.[23,24] Increasing the curvature of membrane is found to increase the magnitude of fluctuation of the GG hinge and the overall tilt of the TMD.[25] It is likely that magnitude of fluctuations in the hinge may enhance Aβ_42_ and Aβ_43_ production.[8] Additionally, simulation studies have revealed[20,26,27] that the GG hinge is an important structural feature for C99 dimers, with the angle of the hinge varying for several distinct dimerization motifs. It has further been noted that the membrane thickness can preferentially stabilize and environmentally select specific C99 dimer strucutres.[17,26–28] Beyond the hinge lies G_38_xxxA_42_, another glycine zipper motif often found in TM dimers,[18] important for C99 homodimerization.[29] The GxxxG repeat motif appears to facilitate C99 dimer formation in thicker membranes while the competing GxxxA motif supports dimers are most often observed in thinner membrane and micelle.[30] At the C-terminal end of the TMD, residues A_42_, T_43_, V_44_, I_45_, V_46_, T_48_, L_52_, and K_53_ all feature several mutations found in AD.[31] Some mutations decrease the propensity for homodimerization,[32] and enhance Aβ_42_ production.[33] A “lysine anchor” formed by the tripe repeat K_53_K_54_K_55_ is evidenced to register at the C-terminal end of the TMD membrane surface.[34]

While the TMD structure has been the focus of experimental and computational studies, the structure of the extra-membrane residues has received relatively little attention in spite of the suspicion that the extra-membrane domains of C99 have interesting features that are crucial to AD. The N-terminus of C99 almost certainly interacts with the nicastrin domain of γ-secretase.[35] Within the N-Loop domain, Ala point mutation of K_28_ has a dramatic impact on APP processing, switching formation of Aβ_40_ to Aβ_33_, implicating this turn in the γ-secretase interaction.[34] Residues 15 to 21 (LVFFAED of the N-Helix domain), sometimes referred to as the juxta-membrane (JM) region, is plays a role in inhibiting γ-secretase binding[36] and binding with cholesterol.[8,13,15] Furthermore, membrane insertion of residues in the JM region appears to sensitively depend on pH.[13] The N-Helix also features mutants A21G,[37] E22Q,[38] E22K,[39] E22G,[40] E22Δ,[41] and D23N,[42] all found to occur in AD patients. Within the NTD, the mutation K16N is known to make APP untenable for binding by α-secretase[43,44] and the E11K mutation was found to enhance Aβ production.[45] The mutations D7H,[46] D7N,[47] H6R,[47] and A2V[48] were found in patients with early onset of AD, suggesting a role for these residues in interaction with γ-secretase. Additionally, histidine residues in the N-terminus H_6_, H13, and H14 are known to bind with Cu and Zn metals, found in high concentration in amyloid plaques.[49] Additionally, Aβ_42_ forms a complex with the C99 N-terminus when C99 is membrane-bound, which enhances C99 homo-oligomer formation.[50]

In the C-Loop there are several phosphorylatable residues, identified at T_58_, S_59_, T_72_,[46] and Y_86_.[51] The phosphorylation of S_59_ enhances trafficking of APP to the golgi apparatus.[52] It has been noted that Ala point mutation at T_72_ may enhance the production of Aβ_40_ and Aβ_42_,[53–55] impacting interaction of APP with some enzymes.[56] Y_86_ has been identified to be phosphorylated at higher concentrations in the brains of AD patients, and is suspected to prevent the interaction of APP with adaptor proteins.[54]

The C-Loop and C-Helix are known to interact with several proteins in the cytoplasm, forming complexes in which these domains adopt an a-helical structure.[57] The C99 sequence binds to many cytoplasmic proteins including the G protein G0 with residues H_61_-K_80_,[58] the adaptor protein Fe65 with residues D_68_-N_99_,[59] the adaptor protein X11 with residues Q_83_-Q_96_,[60] the adaptor protein mDab1 with a similar residues to X11,[61] and the kinase Jip-1with residues N_84_-F_93_.[62] The C-terminus is cleaved by caspases at D_68_ to form C31, a cytoplasmic protein found in AD patients and evidenced to signal apoptosis.[63] Aβ-C99 complex-enhanced C99 oligomerization increases the production of C31.[50] Mutation of D_68_ to Ala prevents production of C31, abrogating cytotoxic function.[64] Additionally, residues 85-91 (GYENPTY) are found to be essential for cytotoxic activity of C31, and are involved in interactions with many cytoplasmic proteins.[50]

Currently, the experimental knowledge of extra-membrane residues of C99 is limited to backbone chemical shifts, NOEs, and EPR signals from membrane-mimicking environments, for which the structural ensembles of residues 6, 12-16, 53-56, 62, 73-76, 80, 81, and 88 are unresolved or too uncertain.[8,9] A prodigious body of work characterizing structure of Aβ fragments has been performed and generally suggests that residues 21-28 of Aβ act as a seed for oligomerization and fibril formation. It has been conjectured that this region contains key residues characterizing the aggregation-prone *N*^*^ state of Aβ and Aβ fragments.[65,66] Support for this conjecture has been provided by NMR and computational studies of Aβ_40_ and Aβ_42_ structural ensembles.[67] Additionally, the N-Turn and C-Loop domains show nearly random coil chemical shifts, implying that they are unstructured on average. However, these domains may exhibit heterogeneity of metastable structural states as has been observed in many intrinsically disordered proteins. The current knowledge of C99 structure and residue features in typical thermodynamic conditions is summarized in Figure 1.

To address some of the outstanding questions related to the structure of C99 and its interaction with the membrane we performed simulations of monomeric wildtype C99 in model membranes, using a computational approach that proved to be remarkably useful in elucidating structures of the TMD.[19,20,27] We employed replica-exchange molecular dynamics (REMD)[68] to sample hundreds of nanoseconds of C99 dynamics at physiological temperatures in 30, 35, and 40 Å-thick membranes modeled using the GBSW implicit solvation method.[69]

In thin membranes we observed extracellular domain states to be correlated with the TMD state. The mean and variance of the TMD and GG hinge angles (Figure 1) were observed to increase with thinning of the membrane. The C-and N-terminal secondary and tertiary structures of C99 were heterogeneous with discrete metastable states. These ensembles were directly compared and contrasted with the results of prior solution NMR and EPR studies. C99 ensembles were found to exhibit newly-observed metastable α-helical and β-strand structures in C- and N-termini, which are correlated with the state of the TMD only in thin membranes. β-strand structures observed in some N-terminal residues are suggestive of templates that may seed amyloid oligomerization on the membrane surface. α-helical domains in the N-terminus are observed and found to be suggestive of nicastrin association sites. α-helical domains observed in previously uncharacterized phosphorylatable sites T_58_, S_59_, and Y_86_ (see Figure 1), suggest that these domains may be involved in interactions that enhance phosphorylation processes. Overall, our work provides an enhanced picture of the structure of extra-membrane residues of C99, and lays the foundation for further investigations of considering the role of C99 structure in facilitating interaction with other molecules in membrane.

## Methods

### A. Initial Structure Preparation

We constructed an initial structure of the full-length C99 sequence using current literature data. Residues 23-55 were modelled using a “Gly-in” structure of one C99 sampled in the recent work by Dominguez et. al.[27] Onto this fragment, residues 1-22 and 56-99 were built using dihedral angles predicted via the TALOS+,[70] using the Cα, Cβ, C, N, and H chemical shifts reported for C99 in LMPG micelles.[30] To remove clashes and effectively move the C-Helix close to the membrane surface, the ψ angle of H_14_, located in the disordered loop of the N-teminus (see Figure 1), was adjusted to 180° and the φ angle of Q_82_ was adjusted to 180°. The rotomeric states of residues 1-22 and 56-99 were assigned using the Shapovalov and Dunbrack rotamer library.[71] Protonation states were assigned using the AddH program in UCSF Chimera,[72] assigning negative GLU and ASP, positive LYS and ARG, neutral CYS and TYR, and setting HIS to the HSD CHARMM histidine type. The center of the membrane was initially set at the z-coordinate of the Cα of G_38_.

### B. Molecular Dynamics Simulation

All simulations were performed using CHARMM version c41b1[73] using the CHARMM36 force field,[74] likely to be the most accurate force field for simulation of Aβ.[75] The GBSW implicit membrane solvent model was used,[69] employing a 0.004 kcal mol^-1^ Å^-2^ surface tension, 5 Å smoothing length from the membrane core-surface boundary, and 0.6 Å smoothing length at the water-membrane surface boundary, using 24 radial and 38 angular integration points to 20 Å. No cutoffs were used for nonbonded interactions. After C99 was inserted in 30, 35, and 40 Å-thick implicit membranes the potential energy was minimized using the steepest descent algorithm until apparent convergence, and simulated for 130, 160, and 460 ns, respectively, using REMD[68] via the REPDSTR utility in CHARMM. We used 16 replicas in REMD simulations, employing exponentially-spaced temperatures from 310 to 500 K and attempting to exchange temperature conditions every 1 ps, manifesting an overall exchange success rate of 17.7 ± 3.7%. Langevin dynamics was employed using a 2 fs time step with a leap frog integrator, a 5 ps^-1^ friction constant, and constrained hydrogen bonds via the SHAKE algorithm. Atomic coordinates were written every 10 ps and all analyses employed coordinate data at this resolution.

### C. Analysis Methods

SciPy,[76,77] Cython,[78] Matplotlib,[79] VMD v1.9.3,[80] MDAnalysis v.0.16.2,[81,82] MDtraj v1.9.0[83] were employed for analyses. SHIFTX2 was used to compute the full set of chemical shifts of C99 in each frame for thermodynamic conditions of 7.4 pH and 310 K temperature.[84] All analyses considered structures sampled at equilibrium (past 30 ns) in the 310 K ensemble from REMD. To assign conformational states of extra-membrane domains of C99, a relatively low-dimensional space that enables precise clustering was constructed. Secondary structure assignments were made using the STRIDE implementation in VMD.

To cluster C99 structures four Principal Component Analysis (PCA) eigenspaces were constructed, using Ca positions and the sine and cosine transformations of dihedral angles (dPCA[85]) of N-terminal residues 1-29 and C-terminal residues 52-99, using data from the equilibrium ensemble in 30, 35, and 40 Å membranes. The first 3 principal components of simulation data in each of these four eigenspaces were considered relevant, each conformation of C99 described by a 12-dimensional space capturing the secondary and tertiary structure of the N-and C-terminus. Conformations at each membrane thickness were assigned to states by clustering in this 12-dimensional space using a 16-cluster Gaussian Mixture Model (GMM). The GMM of simulations at each membrane thickness was constructed using *k*-means clustering to parameterize initial clustering and weights of each data point in each cluster, then refined using 100 iterations of the GMM Expectation-Maximization algorithm.[86] Metric multidimensional scaling of the 12-dimensional data to 2 dimensions for each membrane thickness was performed in order to visualize the nature of the clustering and the relative distance between states.

## Results and Discussion

### A. Convergence of Ensemble to Equilibrium and Experiment

Full C99 sequence was simulated using REMD in GBSW implicit membranes, a successful approach for enhanced sampling of membrane protein structure.[87] Membranes of 30, 35, and 40 Å thicknesses, corresponding to lipids such as DLPC, DMPC, and POPC, were used to study the effect of membrane structure on the conformational ensemble of C99.[88] The initial structure of C99, constructed from a combination of past simulations and chemical shift-based dihedral assignments (Figure 1), gradually evolved in REMD simulations to interact with the membrane surface. The radius of gyration (*R*_*g*_) rapidly converged to the ensemble average in 35 and 40 Å membranes, but appeared to require 20 ns to converge in 30 Å membranes due to relatively slow re-arrangements in secondary structure near the membrane surface (Figure 2A), evidenced by deep insertion of C99 side chains to the membrane (Figure S1). We considered the equilibrium ensemble to have been reached by 30 ns in all REMD simulations, and only consider data at equilibrium for characterization of C99 structure.

**Figure 2.**
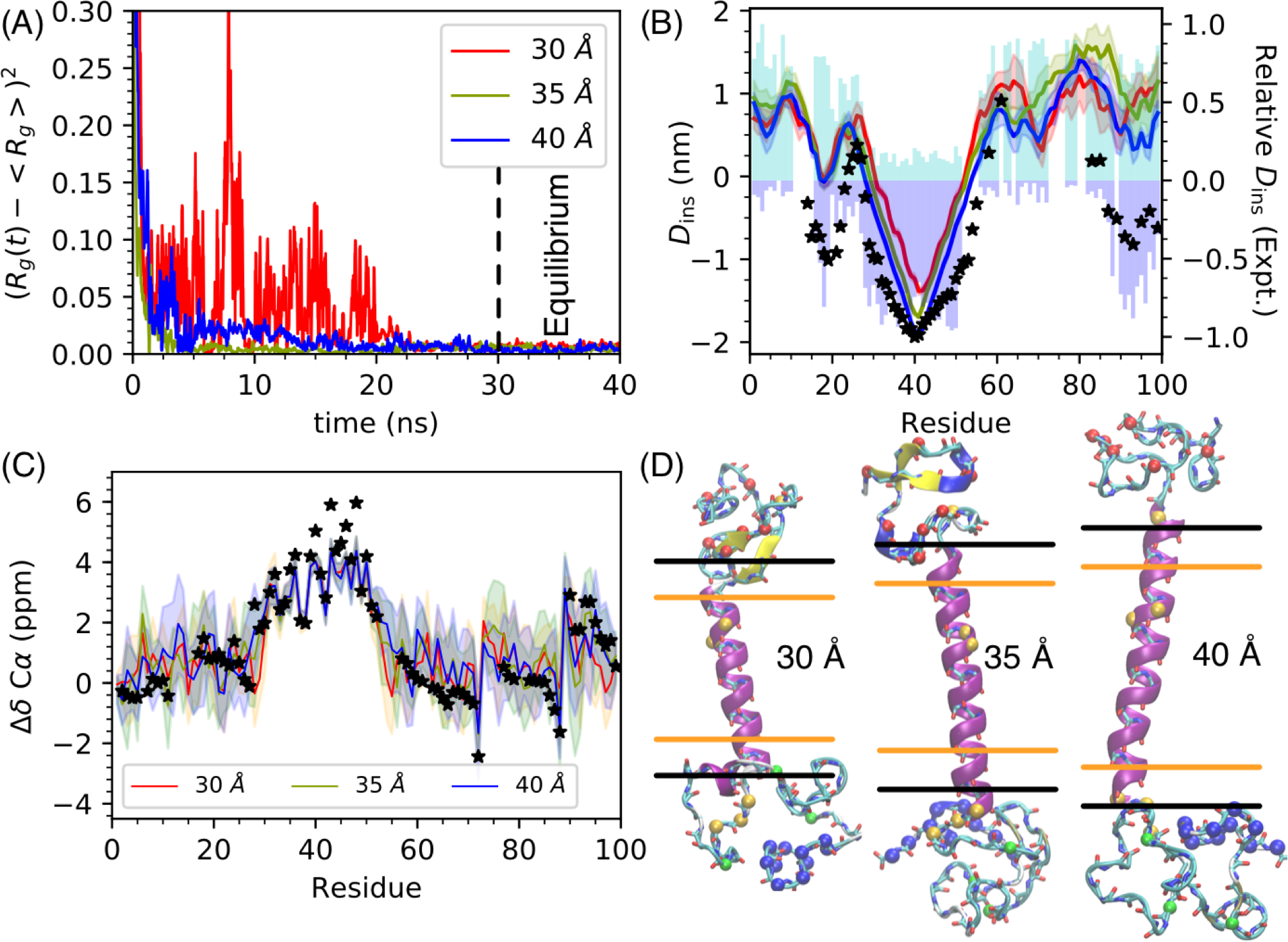
(A) Squared difference of *Rg* from ensemble average over time. The vertical dashed line at 30 ns demarcates the time beyond which the ensembles are considered to be at equilibrium. (B) Equilibrium average and standard deviation of insertion depth of residue Cα in the membrane, *D*_ins_. Stars indicate scaled relative depths of residue insertion to the membrane inferred from EPR probe signals in POPG:POPC membranes.[8] Scaled NMR signals from lipophilic (blue) and hydrophilic (cyan) probes in POPC-DHPC bicelles shown in bars.[9] (C) Equilibrium average and standard deviation of Ca chemical shifts predicted using SHIFTX2. Stars indicate the Ca chemical shifts measured in LMPG micelles. (D) Structures of C99 at 30 ns in 30, 35, and 40 Å membranes. (Same residue coloring as in Figure 1).

The ensemble average of Cα residue depths of insertion (*D*_ins_) in the membrane were well-captured by simulation, comparing well with NMR signals from hydrophobic and hydrophilic probes in POPC-DHPC bicelles and correlating with past EPR measurements in POPG:POPC membranes by Pearson’s *r* of 0.888, 0.861, and 0.894 for 30, 35, and 40 Å membranes, respectively (Figure 2B). This marginally higher insertion depth correlation observed in 40 Å membrane may be attributed to the insertion of the C-Helix in the membrane surface. The whole sequence of the C-Helix was observed to rest on the membrane surface in much of the 40 Å ensemble in contrast with the 30 and 35 Å ensembles that predominantly show residues around T_90_ to rest on the membrane surface. The higher correlation of C99 residue insertion in 40 Å implicit membranes is coincident with POPC membrane, which has been measured to be approximately 40 Å thick in combined analysis of small-angle neutron and X-ray scattering data.[88]

Cα chemical shifts predicted using the SHIFTX2 algorithm, which boasts the best correlations of predicted chemical shifts to experiment of current chemical shift prediction methods, show substantial correlation with those measured in LMPG micelles (*r* correlation coefficients of 0.882, 0.908, and 0.903 for 30, 35, and 40 Å membranes) (Figure 2C). However, overall the 40 Å membrane simulations showed higher correlation with all backbone chemical shifts (Figure S1 and Table S1).[30]

### B. TMD tilt and kink angles

The hinge located at G_37_G_38_ has been conjectured to modify the interaction of C99 with γ-secretase in a way that impacts C99 processing.[21] As the membrane thickness increases the production of Aβ has been reported to increase overall, but the ratio of Aβ_42_/Aβ_40_ decreases.[23,24] This suggests that C99 structures in thick membranes are preferable for appropriate interactions of C99 and γ-secretase. The stability of the TMD helix at the GG hinge has been observed to be weaker than the rest of the TMD via H-D exchange experiments.[21,22]

In past simulation studies the GG hinge flexibility did not appear to be sensitive to membrane thickness.[27] However, the simulations presented here include the full C99 sequence, which seems to be important for sampling certain TMD structures (Figure 3). Here, we define the TMD tilt angle (*θ*) as the angle between the vector of best fit through residue 30-52 Cα positions and the z-axis (Figure 1). The GG hinge angle (*κ*) is the angle between the vectors of best fit through Ca positions of residues 30-37 and of residues 38-52 (Figure 1). In 40 Å membranes there is a single macrostate of TMD structure with average and standard deviation in TMD angles (<*θ*>) of 7.5° ± 3.9° and GG hinge angles (<*κ*>) of 9.3° ± 4.9°. In 35 Å membranes these angles increase to <*θ*> = 11.1° ± 5.6° and <*κ*> = 9.9° ± 5.1°. In 30 Å membranes three structural macrostates of the TMD manifest, composing 29.8% (TM1), 66.2% (TM2), and 4.0% (TM3) of the ensemble. Extramembrane clusters 4, 5, 7, and 8 comprise TM1, featuring <*θ*> of 9.1° ± 3.6° and <*κ*> of 16.6° ± 7.0°. Clusters 1, 2, 3, 6, 9, 10, 12, 13, 14, 15, and 16 comprise TM2, which exhibits <*θ*> of 24.3° ±4.0° and <*κ*> of 14.5° ± 7.0°. Cluster 10 comprises TM3, characterized by <*θ*> of 9.3° ±4.3° and <*κ*> of 46.9° ± 4.8°. The extreme kink in TM3 is an artifact, resulting from unraveling of TMD residues 31-33 to form a β-strand with residues 20-22. These observations suggest that the mean and variance of TMD and GG hinge angles generally increase as a result of membrane thinning. The increase is accompanied by considerable heterogeneity in the C99 structures.

**Figure 3.**
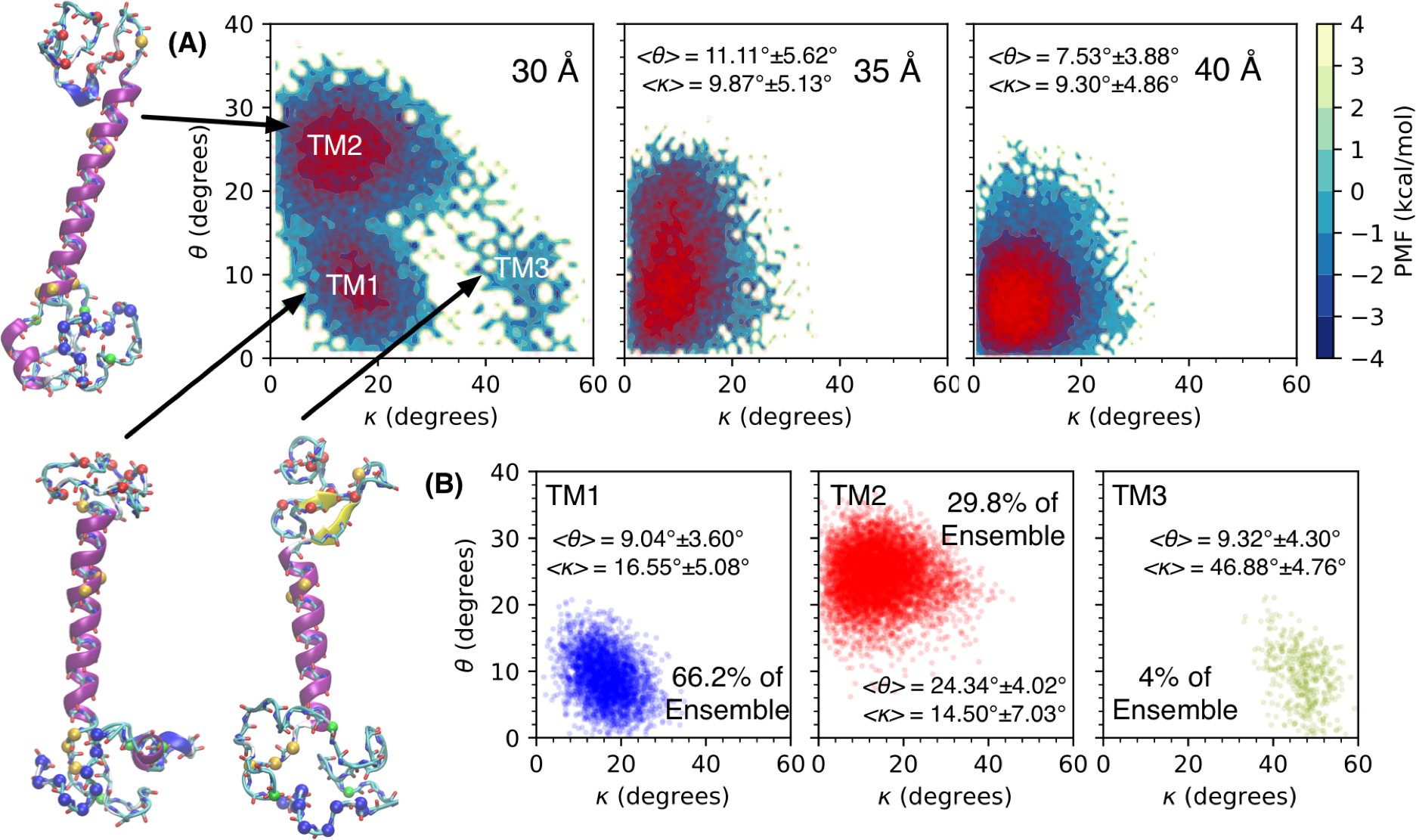
TMD (*θ*) and GG kink *κ* angles of C99 (A) PMF (*-k*_*B*_T ln(*p*)) at equilibrium in 30, 35, and 40 Å membranes, showing 10,000 randomly selected data points in red and (B) 30 Å membrane for TMD macrostates TM1 (blue), TM2 (red), and TM3 (gold), which compose 66.2, 29.8, and 4% of the equilibrium ensemble, respectively. Insets show mean and standard deviation of angles in the displayed macrostate. Representative C99 conformation secondary structure drawn with STRIDE and Ca colored as defined in Figure 1.

### C. Secondary Structure, Membrane Insertion, and Implications of C99 States

The secondary and tertiary structures of extra-membrane residues are heterogeneous. Using projection of simulation data onto a 12-dimensional space the describing relevant PCA eigenvectors of secondary and tertiary structures of N-and C-terminal extra-membrane residues, conformational clusters were assigned and refined using k-means and a Gaussian Mixture Model to find the 16 conformational states defined in 30, 35, and 40 Å membranes (Figs S3-5). These clusters were inspected by embedding the 12-dimensional space to a 2-dimensional space by metric multidimensional scaling and viewing all assigned atomistic structures (Figs S6-8).

Considering the 8 most populated clusters of each membrane, which account for 75.4, 75.4, and 73.0% of 30, 35, and 40 Å membrane ensembles, we identify the most prominent C99 states. Secondary structure assignment via STRIDE allows for the general classification of structure. We consider the secondary structure propensity by taking the difference between the observed α-helix likelihood (*p_α_*) and the observed β-strand likelihood (*p_β_*) (*p*_*s*_ = *p*_*α*_ - *p*_*β*_) for each cluster (Figure 4). In each membrane condition, we observe unique secondary structures including or proximal to sites of non-TMD familial AD mutations, phosphorylatable sites, and the metal binding sites. To consider tertiary structure we measured the insertion depth of Cα to the membrane surface (*D*_ins_).

**Figure 4.**
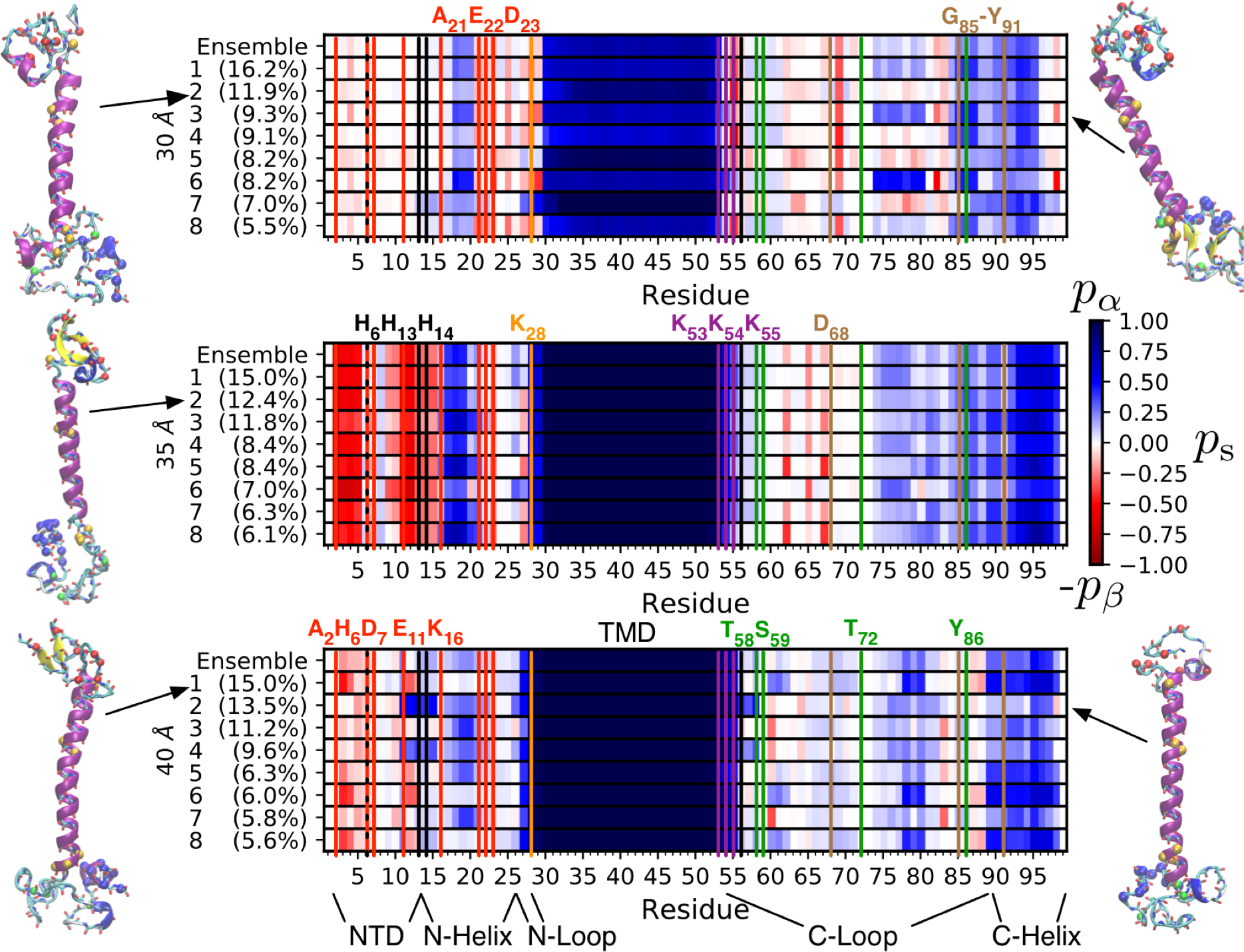
Difference of α and β propensity at each C99 residue in the ensemble (see scale for secondary structure propensity on the right) and in the 8 most populous clusters in 30, 35, and 40 Å membranes (percentages correspond to population of the ensemble). Lines and text indicate residue indices of interest: AD-associated mutations (red), phosphorylation sites (green), lysine anchor (purple), metal binding sites (black), Aβ_33_-producing mutation (orange), and C31 cleavage and cytotoxic function sites (brown). Last frame of visualized C99 clusters with secondary structure drawn with STRIDE and Cα shown as in Figure 1.

The TMD is observed to lengthen on both the C- and N-terminal ends with increasing membrane thickness. The TMD was extended by two residues at the N-terminus and one residue at the C-terminus every 5 Å increase in membrane thickness, growing from residues 30-53 in 30 Å membranes to residues 26-55 in 40 Å membranes. This observation is in contrast to the usual assumption[34] that the lysine anchor does not change its registration with the membrane surface, and that only the N-terminal end of the TMD changes registration with the membrane surface as membrane thickness changes. This had previously been unconfirmed, as past experiments on full-length C99 could not resolve structure or membrane insertion of the lysine anchor in a variety of environments.[9] Residue K_28_, found to change production of Aβ_40_ to Aβ_33_ when mutated to Ala,[34] is incorporated in the TMD helix, undergoing a transition from β to α structure as membrane thickness increases.

In all membrane conditions residues 16-20 have helical propensity and to be inserted to the membrane, in agreement with prior EPR and NMR experiments.[34] The full C-Helix, identified as being inserted in membrane in past experiments, is found to be helical in all conditions other than 30 Å, for which residues 96-99 are unstructured and unassociated with the membrane surface. The C-Helix is observed to be helical even when unassociated with the membrane surface, a condition observed in some clusters in all membrane conditions. This finding provides a structural basis for the conjecture that the C-Helix is available for binding with cytoplasmic proteins in any membrane condition. Residues 73-76, for which membrane insertion and chemical shifts had been previously unresolved in experiments, appear to be unstructured in all membranes and broadly distributed relative to the membrane surface. Residues 74 to 80 are found to be slightly less helical and more bound to the membrane surface in 30 Å membranes, suggesting that thinner membranes may make C99 less available for binding to the G0 protein, which binds residues 61-80.[58] Residue D_68_, the cleavage site for cytotoxic C31 peptide formation, gains more β-propensity as membrane thins. The cytotoxic functional domain G_85_-Y_91_ becomes more a-helical in response to membrane thinning, though the insertion depth does not follow a trend, being membrane-associated in 30 and 40 Å, and membrane-disassociated in 35 Å membranes. It may be possible that C99 in thinner membranes is more amenable to cleavage of D_68_ to form C31.

In 30 Å membranes, residues 21-23, 25-27, and 28-30 occasionally interact to form β-strands, suggestive of the aggregation-prone *N*^*^ structural motif observed in Aβ fragments.[65,66] This structure is not present in 35 and 40 Å membranes, in which residues 28-30 join the TMD helix. Mutants of residues 21-23 are featured in cases of familial AD and thin membranes are known to cause an increase in the ratio of Aβ_42_/Aβ_40_ produced. It is possible that mutations in residues 21-23 stabilize this β-strand, altering the TMD ensemble to resemble the structure observed in 30 Å membranes. Additionally, in some clusters, residue K_55_ forms H-bonds consistent with β-strand structures involving A_69_, occasionally including Q_82_ and Q_83_ as well.

**Figure 5.**
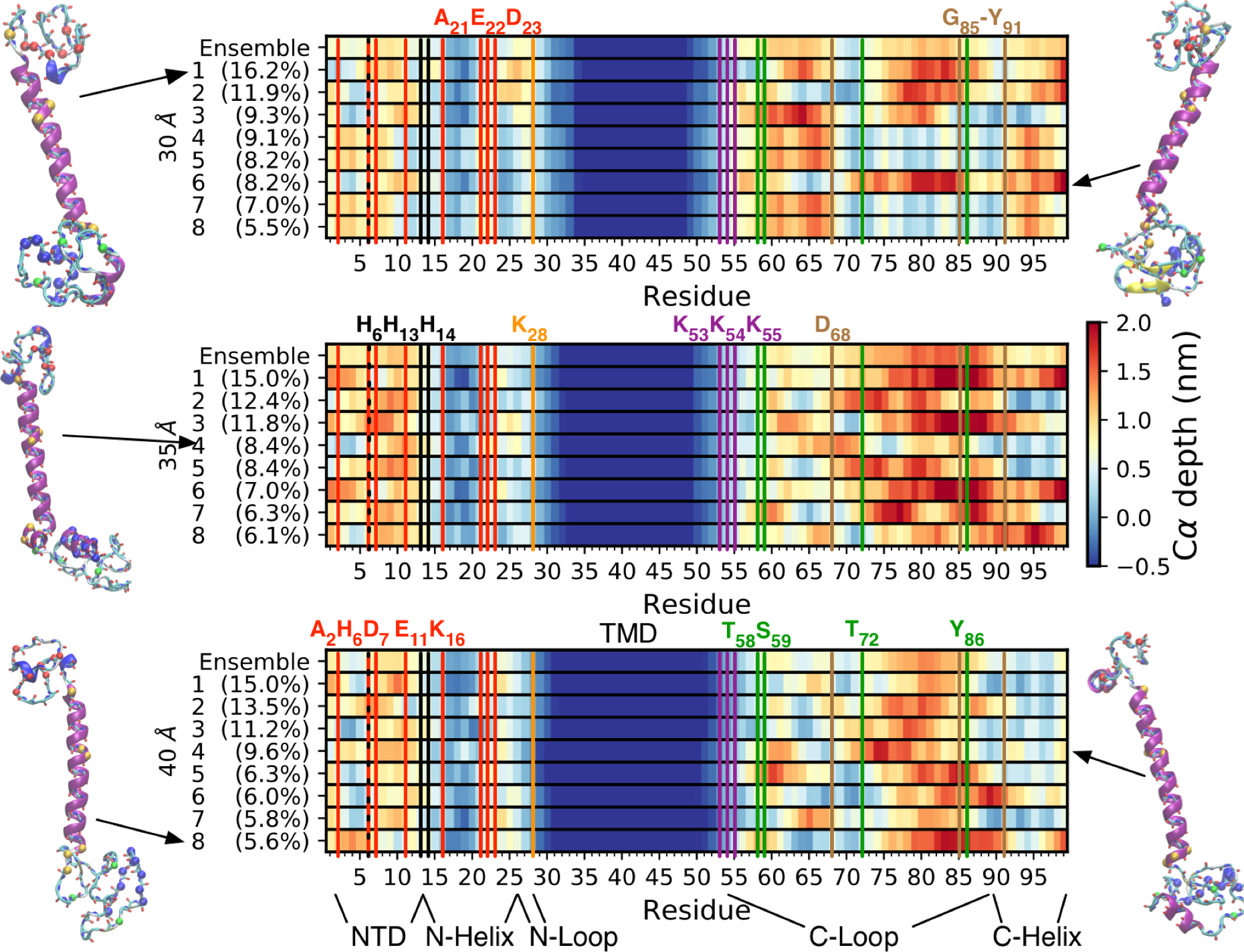
Average membrane insertion of each C99 residue Cα ensemble (see scale for depth of insertion on the right) in the ensemble and in the 8 most populous clusters in 30, 35, and 40 Å membranes (percentages correspond to population of the ensemble). Lines and text indicate residue indices of interest: AD-associated mutations (red), phosphorylation sites (green), lysine anchor (purple), metal binding sites (black), Aβ_33_-producing mutation (orange), and C31 cleavage and cytotoxic function sites (brown). Last frame of indicated C99 clusters with secondary structure drawn with STRIDE and Cα shown as in Figure 1.

In 35 Å membranes, a prominent β-hairpin is formed with residues 2-5 and 11-15, in which residue N_27_ sometimes participates via H-bonding. This hairpin is positioned away from the membrane surface. This structure does not appear in membrane-bound α_1-42_ in similar implicit membrane simulations,[20] and seems to be unique to 35 Å-thick membranes. It is possible that this β-hairpin structure acts as a seed for Aβ oligomerization. C99-seeded Aβ association to the membrane may be much more favorable than pure Aβ mixtures considered in the past,[89] as Aβ is at substantially higher concentration outside the cells than in the membrane. Mutation of residues 2, 11, and 16, found in familial AD, may change the propensity for this β-hairpin to form. Additionally, H_6_, H_13_, and H_14_, residues known to bind metal ions found at high concentration in amyloid plaques,[49] are proximate to the observed β-hairpin. The structure of His-ion complexes found in computational investigations of Aβ_1-16_ resembles this hairpin structure.[90,91] As such it may be possible that 35 Å membranes are ideal for stabilizing C99 structures that bind metal ions.

In 40 Å membranes there is weaker propensity for β-hairpin formation observed in 35 Å membranes. A strand with residues 2-5 and 11-13 in some clusters, such as 1, 6, and 8, is observed. In clusters 2 and 4, residues 11-15 form an α-helix that is unassociated with the membrane. Along with residues 16-20, this helix may serve as an interaction site with the nicastrin domain of γ-secretase. The formation of these α-helices may serve to enhance the recognition of C99 by γ-secretase as one possible mechanism explaining the observed increase in Aβ processing observed in thicker membranes.

**Figure 6.**
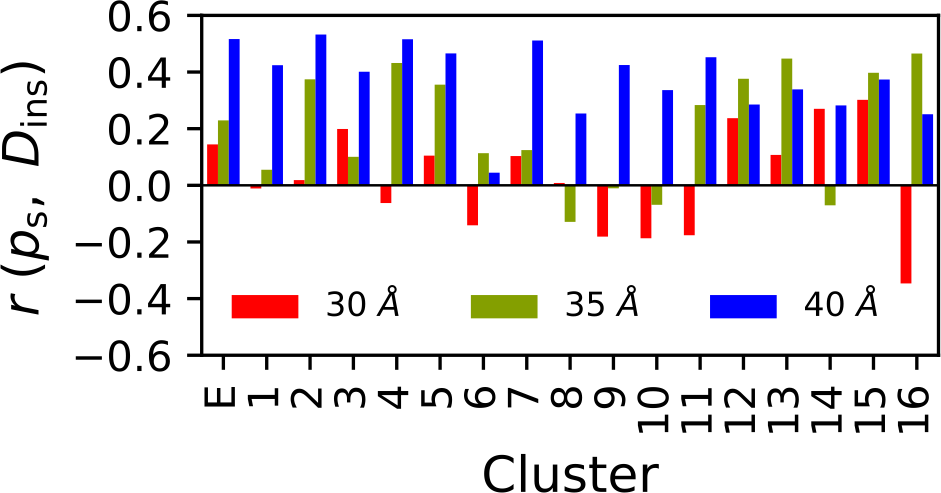
Pearson’s *r* correlation of average Cα depth of insertion in membrane (*D*_ins_) and difference in observed α and β structure propensity (*p_s_*) in C99 residues 1-28 and 53-99 in the equilibrium ensemble (E) and in each cluster.

The secondary structure propensities and insertion depth of non-TMD residues 1-28 and 52-99 for the whole ensemble and for each cluster reveal that the helicity of extra-membrane residues is not correlated with membrane insertion depth. This observation is contrary to typical expectation that the more hydrophobic membrane environment increases the propensity for helical structure. This is quantified by Pearson’s *r* correlation of non-TMD residue secondary structure propensity to membrane insertion of Cα, *r*(*p*_*s*_, *D*_ins_), in the ensemble and in clusters (Figure 6). It is indeed possible that this could be a consequence of the simulation model used, and further investigation using explicit solvent simulations with consideration of the disordered protein structure should be pursued.

## Conclusions

We performed REMD simulations of full length C99 in model membranes of 30, 35, and 40 Å thicknesses. We observe the TMD and G_37_G_38_ hinge angle means and variances to increase as the membrane thins. Multiple TMD states were found to manifested in thin membranes, with which a positive correlation between TMD states with extra-membrane residue states was found. Heterogeneous but discrete structural states observed in the C99 C- and N-terminal extramembrane regions of C99 are found to be unique to the specific membrane condition. Generally, an increase in α and β secondary structure is observed as membrane thickness increases. The TMD helix expands on the N- and C-terminal ends as membrane thickness increases. Residues 21-23, 25-27, and 28-30 form β-strands similar to the aggregation-prone *N*^*^ motif previously observed in Aβ fragments in 30 Å membranes. In addition, residues 2-5 and 11-15 form a β-hairpin in 35 Å membranes. It is conjectured that these β-strand motifs may serve as seeds for Aβ aggregation on the membrane surface. Residues 11-15 adopt α-helical structures in 40 Å membranes that may promote binding of C99 with the nicastrin domain of γ-secretase to promote non-amyloidogenic processing of C99. α-helical structures are generally stabilized in the C-terminus as membrane thickness increases, and do not require association with the membrane surface. This observation suggests that these domains are readily available to interact with proteins in the cytoplasm. Conversely, residues 85-91, known to be essential to cytotoxic function, become α-helical as the membrane thins. These observations drawn from our simulation study are summarized in Figure 7.

**Figure 7.**
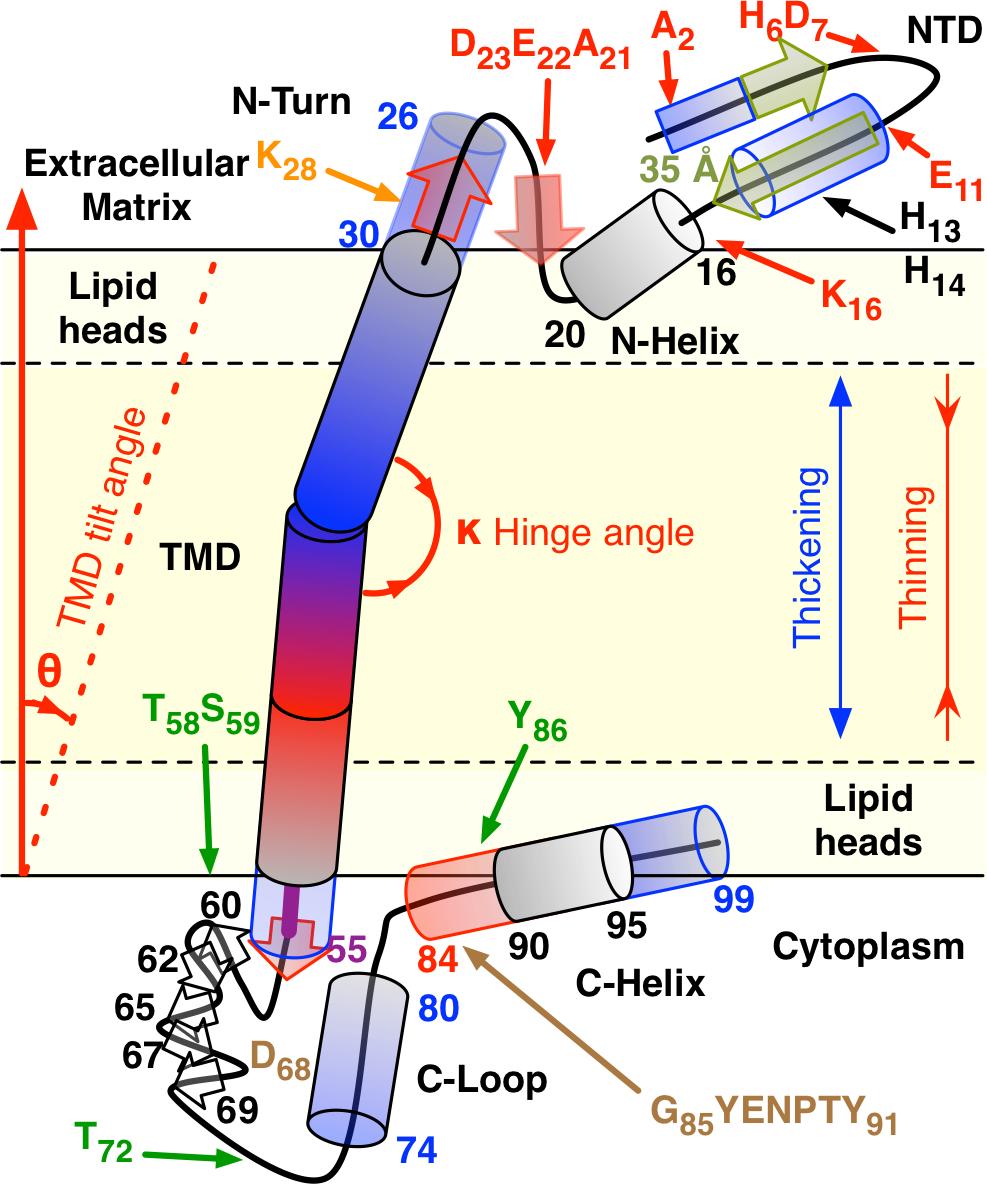
Yellow shading represents the hydrophobic environment of the membrane. Secondary structures resulting from membrane thinning (red), membrane thickening (blue), and unique to 35 Å membranes (gold) are transparent. Residue indices are provided to identify regions in which secondary structure is observed. Red residues are found mutated in familial AD, black residues are important for metal binding, green residues are phosphorylatable, orange K_28_ Ala mutation changes Aβ produced, and brown residues are critical for C31 formation and cytotoxicity. TMD and GG hinge angle means and variance increase with thinning membrane. Black-outlined C-loop β-strands are transient in many membrane conditions.

The insights provided by this study enhance our current understanding of the structural ensemble of full length C99 in membrane and the potential role played by C99 structure in recognition and processing by γ-secretase. Taken together, these results open the path to investigations of the role of C99 structure in interactions with γ-secretase and Aβ, which may lead to new perspectives on the genesis of amyloid in AD.

## Acknowledgements

G.A.P. thanks the NSF GRFP for support under NSF Grant No. DGE-1247312, the NSF GROW program, and the 2017 JSPS Postdoctoral Fellowships for Research in Japan Strategic Program. J. E. S. and D. T. acknowledge the generous support of the National Institutes of Health (R01 GM107703). D.T. thanks the Collie-Welch Regents chair (F0019) for generous support. We thank Afra Panahi for discussion regarding phosphorylatable sites on C99 and assistance in preparation of simulations using the REPDSTR utility. The authors acknowledge the Shared Computing Cluster, administered by Boston University’s Research Computing Services, used for many of the MD simulations.

## References

[1] C. Priller, T. Bauer, G. Mitteregger, B. Krebs, H.A. Kretzschmar, J. Herms, Synapse Formation and Function Is Modulated by the Amyloid Precursor Protein, J. Neurosci. 26 (2006) 7212–7221. doi:10.1523/JNEUROSCI.1450-06.2006.

[2] J. Morales-Corraliza, M.J. Mazzella, J.D. Berger, N.S. Diaz, J.H.K. Choi, E. Levy, Y. Matsuoka, E. Planel, P.M. Mathews, In Vivo Turnover of Tau and APP Metabolites in the Brains of Wild-Type and Tg2576 Mice: Greater Stability of sAPP in the β-Amyloid Depositing Mice, PLoS One. 4 (2009) e7134. doi:10.1371/journal.pone.0007134.

[3] T. Tomita, Molecular mechanism of intramembrane proteolysis by γ-secretase, J. Biochem. 156 (2014) 195–201. doi:10.1093/jb/mvu049.

[4] U. Sengupta, A.N. Nilson, R. Kayed, The Role of Amyloid-β Oligomers in Toxicity, Propagation, and Immunotherapy, EBioMedicine. 6 (2016) 42–49. doi:10.1016/j.ebiom.2016.03.035.

[5] A. Prasansuklab, T. Tewin, Amyloidosis in Alzheimer’s Disease: The Toxicity of Amyloid Beta (A, Evidence-Based Complement. Altern. Med. 2013 (2013) 10 pages. doi:10.1155.

[6] R. Kayed, Common Structure of Soluble Amyloid Oligomers Implies Common Mechanism of Pathogenesis, Science (80-.). 300 (2003) 486–489. doi:10.1126/science.1079469.

[7] G.M. Shankar, S. Li, T.H. Mehta, A. Garcia-munoz, E. Nina, I. Smith, F.M. Brett, M. a Farrell, M.J. Rowan, C. a Lemere, C.M. Regan, D.M. Walsh, B.L. Sabatini, D.J. Selkoe, Amyloid-beta protein dimers isloated directly from Alzheimer brain impair synaptic plasticity and memory, Nat. Med. 14 (2008) 837–842. doi:10.1038/nm1782.Amyloid.

[8] P. J. Barrett, Y. Song, W.D. Van Horn, E.J. Hustedt, J.M. Schafer, A. Hadziselimovic, A. J. Beel, C.R. Sanders, The Amyloid Precursor Protein Has a Flexible Transmembrane Domain and Binds Cholesterol, Science (80-.). 336 (2012) 1168–1171. 21 doi:10.1126/science.1219988.

[9] Y. Song, K.F. Mittendorf, Z. Lu, C.R. Sanders, Impact of Bilayer Lipid Composition on the Structure and Topology of the Transmembrane Amyloid Precursor C99 Protein, J. Am. Chem. Soc. 136 (2014)4093–4096. doi:10.1021/ja4114374.

[10] T. Tomita, T. Iwatsubo, Structural biology of presenilins and signal peptide peptidases, J. Biol. Chem. 288 (2013) 14673–14680. doi:10.1074/jbc.R113.463281.

[11] M.M. Javadpour, M. Eilers, M. Groesbeek, S.O. Smith, Helix packing in polytopic membrane proteins: Role of glycine in transmembrane helix association, Biophys. J. 77 (1999) 1609–1618. doi:10.1016/S0006-3495(99)77009-8.

[12] S. Kim, T.-J. Jeon, A. Oberai, D. Yang, J.J. Schmidt, J.U. Bowie, Transmembrane glycine zippers: Physiological and pathological roles in membrane proteins, Proc. Natl. Acad. Sci. 102 (2005) 14278–14283. doi:10.1073/pnas.0501234102.

[13] A. Panahi, A. Bandara, G.A. Pantelopulos, L. Dominguez, J.E. Straub, Specific binding of cholesterol to C99 domain of amyloid precursor protein depends critically on charge state of protein, J. Phys. Chem. Lett. 7 (2016) 3535–3541. doi:10.1021/acs.jpclett.6b01624.

[14] A.J. Beel, M. Sakakura, P.J. Barrett, C.R. Sanders, Direct binding of cholesterol to the amyloid precursor protein: An important interaction in lipid-Alzheimer’s disease relationships?, Biochim. Biophys. Acta - Mol. Cell Biol. Lipids. 1801 (2010) 975–982. doi:10.1016/j.bbalip.2010.03.008.

[15] Y. Song, E.J. Hustedt, S. Brandon, C.R. Sanders, Competition Between Homodimerization and Cholesterol Binding to the C99 Domain of the Amyloid Precursor Protein, Biochemistry. 52 (2013) 5051–5064. doi:10.1021/bi400735x.

[16] N. Pierrot, D. Tyteca, L. D’auria, I. Dewachter, P. Gailly, A. Hendrickx, B. Tasiaux, L. El Haylani, N. Muls, F. N’Kuli, A. Laquerrière, J.B. Demoulin, D. Campion, J.P. Brion, P.J. Courtoy, P. Kienlen-Campard, J.N. Octave, Amyloid precursor protein controls cholesterol turnover needed for neuronal activity, EMBO Mol. Med. 5 (2013) 608–625. doi:10.1002/emmm.201202215.

[17] C.-D. Li, Q. Xu, R.-X. Gu, J. Qu, D.-Q. Wei, The dynamic binding of cholesterol to the multiple sites of C99: as revealed by coarse-grained and all-atom simulations, Phys. Chem. Chem. Phys. 19 (2017) 3845–3856. doi:10.1039/C6CP07873G.

[18] S.M. Anderson, B.K. Mueller, E.J. Lange, A. Senes, Combination of Cα–H Hydrogen 22 Bonds and van der Waals Packing Modulates the Stability of GxxxG-Mediated Dimers in Membranes, J. Am. Chem. Soc. 139 (2017) 15774–15783. doi:10.1021/jacs.7b07505.

[19] N. Miyashita, J.E. Straub, D. Thirumalai, Y. Sugita, Transmembrane structures of amyloid precursor protein dimer predicted by replica-exchange molecular dynamics simulations, J. Am. Chem. Soc. 131 (2009) 3438–3439. doi:10.1021/ja809227c.

[20] N. Miyashita, J.E. Straub, D. Thirumalai, Structures of β-Amyloid Peptide 1-40, 1-42, and 1-55—the 672-726 Fragment of APP—in a Membrane Environment with Implications for Interactions with γ-Secretase, J. Am. Chem. Soc. 131 (2009) 17843–17852. doi:10.1021/ja905457d.

[21] O. Pester, P.J. Barrett, D. Hornburg, P. Hornburg, R. Pröbstle, S. Widmaier, C. Kutzner, M. Dürrbaum, A. Kapurniotu, C.R. Sanders, C. Scharnagl, D. Langosch, The backbone dynamics of the amyloid precursor protein transmembrane helix provides a rationale for the sequential cleavage mechanism of gamma-secretase, J. Am. Chem. Soc. 135 (2013) 1317–1329. doi:10.1021/ja3112093.

[22] Z. Cao, J.M. Hutchison, C.R. Sanders, J.U. Bowie, Backbone Hydrogen Bond Strengths Can Vary Widely in Transmembrane Helices, J. Am. Chem. Soc. 139 (2017) 10742–10749. doi:10.1021/jacs.7b04819.

[23] E. Winkler, F. Kamp, J. Scheuring, A. Ebke, A. Fukumori, H. Steiner, Generation of Alzheimer disease-associated amyloid beta 42/43 peptide by gamma-secretase can be inhibited directly by modulation of membrane thickness, J. Biol. Chem. 287 (2012) 21326–21334. doi:10.1074/jbc.M112.356659.

[24] O. Holmes, S. Paturi, W. Ye, M.S. Wolfe, D.J. Selkoe, Effects of Membrane Lipids on the Activity and Processivity of Purified γ-Secretase, Biochemistry. 51 (2012) 3565–3575. doi:10.1021/bi300303g.

[25] L. Dominguez, S.C. Meredith, J.E. Straub, D. Thirumalai, Transmembrane Fragment Structures of Amyloid Precursor Protein Depend on Membrane Surface Curvature, J. Am. Chem. Soc. 136 (2014) 854–857. doi:10.1021/ja410958j.

[26] L. Dominguez, L. Foster, S.C. Meredith, J.E. Straub, D. Thirumalai, Structural Heterogeneity in Transmembrane Amyloid Precursor Protein Homodimer Is a Consequence of Environmental Selection, J. Am. Chem. Soc. 136 (2014) 9619–9626. doi:10.1021/ja503150x.23

[27] L. Dominguez, L. Foster, J.E. Straub, D. Thirumalai, Impact of membrane lipid composition on the structure and stability of the transmembrane domain of amyloid precursor protein, Proc. Natl. Acad. Sci. 113 (2016) E5281--E5287. doi:10.1073/pnas.1606482113.

[28] S. Viswanath, L. Dominguez, L.S. Foster, J.E. Straub, R. Elber, Extension of a protein docking algorithm to membranes and applications to amyloid precursor protein dimerization, Proteins Struct. Funct. Bioinforma. 83 (2015) 2170–2185. doi:10.1002/prot.24934.

[29] M. Audagnotto, T. Lemmin, A. Barducci, M. Dal Peraro, Effect of the Synaptic Plasma Membrane on the Stability of the Amyloid Precursor Protein Homodimer, J. Phys. Chem. Lett. 7 (2016) 3572–3578. doi:10.1021/acs.jpclett.6b01721.

[30] A.J. Beel, C.K. Mobley, H.J. Kim, F. Tian, A. Hadziselimovic, B. Jap, J.H. Prestegard, C.R. Sanders, Structural Studies of the Transmembrane C-Terminal Domain of the Amyloid Precursor Protein (APP): Does APP Function as a Cholesterol Sensor?, Biochemistry. 47 (2008) 9428–9446. doi:10.1021/bi800993c.

[31] S. Weggen, D. Beher, Molecular consequences of amyloid precursor protein and presenilin mutations causing autosomal-dominant Alzheimer’s disease., Alzheimers. Res. Ther. 4 (2012) 9. doi:10.1186/alzrt107.

[32] Y. Yan, T.-H. Xu, K.G. Harikumar, L.J. Miller, K. Melcher, H.E. Xu, Dimerization of the transmembrane domain of amyloid precursor protein is determined by residues around the gamma-secretase cleavage sites, J. Biol. Chem. 292 (2017) jbc.M117.789669. doi:10.1074/jbc.M117.789669.

[33] M. Dimitrov, J.R. Alattia, T. Lemmin, R. Lehal, A. Fligier, J. Houacine, I. Hussain, F. Radtke, M. Dal Peraro, D. Beher, P.C. Fraering, Alzheimers disease mutations in APP but not 3-secretase modulators affect epsilon-cleavage-dependent AICD production, Nat. Commun. 4 (2013). doi:10.1038/ncomms3246.

[34] T.L. Kukar, T.B. Ladd, P. Robertson, S.A. Pintchovski, B. Moore, M.A. Bann, Z. Ren, K. Jansen-West, K. Malphrus, S. Eggert, H. Maruyama, B.A. Cottrell, P. Das, G.S. Basi, E.H. Koo, T.E. Golde, Lysine 624 of the Amyloid Precursor Protein (APP) Is a Critical Determinant of Amyloid β Peptide Length, J. Biol. Chem. 286 (2011) 39804–39812. doi:10.1074/jbc.M111.274696.

[35] J.H. Goo, W.J. Park, Elucidation of the Interactions between C99, Presenilin, and Nicastrin by the Split-Ubiquitin Assay, DNA Cell Biol. 23 (2004) 59–65. doi:10.1089/104454904322745934.

[36] Y. Tian, B. Bassit, D. Chau, Y.M. Li, An APP inhibitory domain containing the Flemish mutation residue modulates γ-secretase activity for AB production, Nat. Struct. Mol. Biol. 17 (2010) 151–158. doi:10.1038/nsmb.1743.

[37] L. Hendriks, C.M. van Duijn, P. Cras, M. Cruts, W. Van Hul, F. van Harskamp, A. Warren, M.G. McInnis, S.E. Antonarakis, J.-J. Martin, A. Hofman, C. Van Broeckhoven, Presenile dementia and cerebral haemorrhage linked to a mutation at codon 692 of the β–amyloid precursor protein gene, Nat. Genet. 1 (1992)218–221. doi:10.1038/ng0692-218.

[38] E. Levy, M. Carman, I. Fernandez-Madrid, M. Power, I. Lieberburg, S. van Duinen, G. Bots, W. Luyendijk, B. Frangione, Mutation of the Alzheimer’s disease amyloid gene in hereditary cerebral hemorrhage, Dutch type, Science (80-.). 248 (1990) 1124–1126. doi:10.1126/science.2111584.

[39] O. Bugiani, A. Padovani, M. Magoni, G. Andora, M. Sgarzi, M. Savoiardo, A. Bizzi, G. Giaccone, G. Rossi, F. Tagliavini, An Italian type of HCHWA, Neurobiol Aging. 19 (1998) S238.

[40] C. Nilsberth, A. Westlind-Danielsson, C.B. Eckman, M.M. Condron, K. Axelman, C. Forsell, C. Stenh, J. Luthman, D.B. Teplow, S.G. Younkin, J. Näslund, L. Lannfelt, The “Arctic” APP mutation (E693G) causes Alzheimer’s disease by enhanced Aβ protofibril formation, Nat. Neurosci. 4 (2001) 887–893. doi:10.1038/nn0901-887.

[41] T. Tomiyama, T. Nagata, H. Shimada, R. Teraoka, A. Fukushima, H. Kanemitsu, H. Takuma, R. Kuwano, M. Imagawa, S. Ataka, Y. Wada, E. Yoshioka, T. Nishizaki, Y. Watanabe, H. Mori, A new amyloid beta variant favoring oligomerization in Alzheimer’s-type dementia, Ann. Neurol. 63 (2008) 377–387. doi:10.1002/ana.21321.

[42] T.J. Grabowski, H.S. Cho, J.P.G. Vonsattel, G. William Rebeck, S.M. Greenberg, Novel amyloid precursor protein mutation in an Iowa family with dementia and severe cerebral amyloid angiopathy, Ann. Neurol. 49 (2001) 697–705. doi:10.1002/ana.1009.

[43] M. Citron, T. Oltersdorf, C. Haass, L. McConlogue, a Y. Hung, P. Seubert, C. Vigo-Pelfrey, I. Lieberburg, D.J. Selkoe, Mutation of the beta-amyloid precursor protein in familial Alzheimer’s disease increases beta-protein production., Nature. 360 (1992) 672–674. doi:10.1038/360672a0.

[44] D. Kaden, A. Harmeier, C. Weise, L.M. Munter, V. Althoff, B.R. Rost, P.W. Hildebrand, D. Schmitz, M. Schaefer, R. Lurz, S. Skodda, R. Yamamoto, S. Arlt, U. Finckh, G. Multhaup, Novel APP/Abeta mutation K16N produces highly toxic heteromeric Abeta oligomers, EMBO Mol. Med. 4 (2012) 647–659. doi:10.1002/emmm.201200239.

[45] L. Zhou, N. Brouwers, I. Benilova, A. Vandersteen, M. Mercken, K. Van Laere, P. Van Damme, D. Demedts, F. Van Leuven, K. Sleegers, K. Broersen, C. Van Broeckhoven, R. Vandenberghe, B. De Strooper, Amyloid precursor protein mutation E682K at the alternative β-secretase cleavage β’-site increases Aβ generation, EMBO Mol. Med. 3 (2011) 291–302. doi:10.1002/emmm.201100138.

[46] M. Oishi, A.C. Nairn, A.J. Czernik, G.S. Lim, T. Isohara, S.E. Gandy, P. Greengard, T. Suzuki, The cytoplasmic domain of Alzheimer’s amyloid precursor protein is phosphorylated at Thr654, Ser655, and Thr668 in adult rat brain and cultured cells., Mol. Med. 3 (1997) 111–23. http://www.pubmedcentral.nih.gov/articlerender.fcgi?artid=2230054&tool=pmcentrez&rendertype=abstract.

[47] Y. Wakutani, K. Watanabe, Y. Adachi, K. Wada-Isoe, K. Urakami, H. Ninomiya, T.C. Saido, T. Hashimoto, T. Iwatsubo, K. Nakashima, Novel amyloid precursor protein gene missense mutation (D678N) in probable familial Alzheimer’s disease, J. Neurol. Neurosurg. Psychiatry. 75 (2004) 1039–1042. doi:10.1136/jnnp.2003.010611.

[48] G. Di Fede, M. Catania, M. Morbin, G. Rossi, S. Suardi, G. Mazzoleni, M. Merlin, A.R. Giovagnoli, S. Prioni, A. Erbetta, C. Falcone, M. Gobbi, L. Colombo, A. Bastone, M. Beeg, C. Manzoni, B. Francescucci, A. Spagnoli, L. Cantu, E. Del Favero, E. Levy, M. Salmona, F. Tagliavini, A Recessive Mutation in the APP Gene with Dominant-Negative Effect on Amyloidogenesis, Science (80-.). 323 (2009) 1473–1477. doi:10.1126/science.1168979.

[49] C.J. Maynard, A.I. Bush, C.L. Masters, R. Cappai, Q.-X. Li, Metals and amyloid-beta in Alzheimer’s disease., Int. J. Exp. Pathol. 86 (2005) 147–159. doi:10.1111/j.0959-9673.2005.00434.x.

[50] G.M. Shaked, M.P. Kummer, D.C. Lu, V. Galvan, D.E. Bredesen, E.H. Koo, Abeta induces cell death by direct interaction with its cognate extracellular domain on APP (APP 26 597-624)., FASEB J. 20 (2006) 1254–1246. doi:10.1096/fj.05-5032fje.

[51] N. Zambrano, P. Bruni, G. Minopoli, R. Mosca, D. Molino, C. Russo, G. Schettini, M. Sudol, T. Russo, The β-Amyloid Precursor Protein APP is Tyrosine-Phosphorylated in Cells Expressing a Constitutively Active Form of the Abl Protoncogene, J. Biol. Chem. 276 (2001) 19787–19792. doi:10.1074/jbc.M100792200.

[52] S.I. Vieira, S. Rebelo, S.C. Domingues, E.F. Cruz e Silva, O.A.B. Cruz e Silva, S655 phosphorylation enhances APP secretory traffic, Mol. Cell. Biochem. 328 (2009) 145–154. doi:10.1007/s11010-009-0084-7.

[53] C. Feyt, N. Pierrot, B. Tasiaux, J. Van Hees, P. Kienlen-Campard, P.J. Courtoy, J.N. Octave, Phosphorylation of APP695 at Thr668 decreases gamma-cleavage and extracellular Abeta, Biochem. Biophys. Res. Commun. 357 (2007) 1004–1010. doi:10.1016/j.bbrc.2007.04.036.

[54] E. Poulsen, F. Iannuzzi, H. Rasmussen, T. Maier, J. Enghild, A. Jørgensen, C. Matrone, An aberrant phosphorylation of amyloid precursor protein tyrosine regulates its trafficking and the binding to the Clathrin endocytic complex in neural stem cells of Alzheimer’s disease patients, Front. Mol. Neurosci. 10 (2017) 59. doi:10.3389/fnmol.2017.00059.

[55] Y. Sano, T. Nakaya, S. Pedrini, S. Takeda, K. Iijima-Ando, K. Iijima, P.M. Mathews, S. Itohara, S. Gandy, T. Suzuki, Physiological Mouse Brain Aβ Levels Are Not Related to the Phosphorylation State of Threonine-668 of Alzheimer’s APP, PLoS One. 1 (2006) e51. doi:10.1371/journal.pone.0000051.

[56] T. Suzuki, M. Oishi, D.R. Marshak, A.J. Czernik, A.C. Nairn, P. Greengard, Cell Cycle-Dependent Regulation of the Phosphorylation and Metabolism of the Alzheimer Amyloid Precursor Protein, EMBO J. 13 (1994) 1114–1122.

[57] T.A. Ramelot, L.N. Gentile, L.K. Nicholson, Transient structure of the amyloid precursor protein cytoplasmic tail indicates preordering of structure for binding to cytosolic factors, Biochemistry. 39 (2000) 2714–2725. doi:10.1021/bi992580m.

[58] U. Giambarella, T. Yamatsuji, T. Okamoto, T. Matsui, T. Ikezu, Y. Murayama, M.A. Levine, A. Katz, N. Gautam, I. Nishimoto, G protein beta gamma complex-mediated apoptosis by familial Alzheimer’s disease mutant of APP, EMBO J. 16 (1997) 4897–4907.

[59] T. Russo, R. Faraonio, G. Minopoli, P. De Candia, S. De Renzis, N. Zambrano, Fe65 and 27 the protein network centered around the cytosolic domain of the Alzheimer’s beta-amyloid precursor protein, FEBS Lett. 434 (1998) 1–7. doi:10.1016/S0014-5793(98)00941-7.

[60] Z. Zhang, C.H. Lee, V. Mandiyan, J.P. Borg, B. Margolis, J. Schlessinger, J. Kuriyan, Sequence-specific recognition of the internalization motif of the Alzheimer’s amyloid precursor protein by the X11 PTB domain, EMBO J. 16 (1997) 6141–6150. doi:10.1093/emboj/16.20.6141.

[61] L. Parisiadou, S. Efthimiopoulos, Expression of mDab1 promotes the stability and processing of amyloid precursor protein and this effect is counteracted by X11??, Neurobiol. Aging. 28 (2007) 377–388. doi:10.1016/j.neurobiolaging.2005.12.015.

[62] M.H. Scheinfeld, R. Roncarati, P. Vito, P.A. Lopez, M. Abdallah, L. D’Adamio, Jun NH2-terminal kinase (JNK) interacting protein 1 (JIP1) binds the cytoplasmic domain of the Alzheimer’s β-amyloid precursor protein (APP), J. Biol. Chem. 277 (2002) 3767–3775. doi:10.1074/jbc.M108357200.

[63] D.C. Lu, S. Rabizadeh, S. Chandra, R.F. Shayya, L.M. Ellerby, X. Ye, G.S. Salvesen, E. H. Koo, D.E. Bredesen, A second cytotoxic proteolytic peptide derived from amyloid β-protein precursor, Nat. Med. 6 (2000) 397–404. doi:10.1038/74656.

[64] D.C. Lu, S. Soriano, D.E. Bredesen, E.H. Koo, Caspase cleavage of the amyloid precursor protein modulates amyloid β-protein toxicity, J. Neurochem. 87 (2003) 733–741. doi:10.1046/j.1471-4159.2003.02059.x.

[65] B. Tarus, J.E. Straub, D. Thirumalai, Dynamics of Asp23-Lys28 Salt-Bridge Formation in Abeta 10-35 Monomers, J. Am. Chem. Soc. 128 (2006) 16159–16168. doi:10.1021/ja064872y.

[66] J.E. Straub, D. Thirumalai, Toward a molecular theory of early and late events in monomer to amyloid fibril formation, Annu. Rev. Phys. Chem. 62 (2011) 437–463. doi:10.1146/annurev-physchem-032210-103526.

[67] N. L. Fawzi, A.H. Phillips, J.Z. Ruscio, M. Doucleff, D.E. Wemmer, T. Head-Gordon, Structure and Dynamics of the Aβ 21–30 Peptide from the Interplay of NMR Experiments and Molecular Simulations, J. Am. Chem. Soc. 130 (2008) 6145–6158. doi:10.1021/ja710366c.

[68] Y. Sugita, Y. Okamoto, Replica exchange molecular dynamics method for protein folding 28 simulation., Chem. Phys. Lett. 314 (1999) 141–151. doi:10.1016/S0009-2614(99)01123-9.

[69] W. Im, M. Feig, C.L. Brooks, An Implicit Membrane Generalized Born Theory for the Study of Structure, Stability, and Interactions of Membrane Proteins, Biophys. J. 85 (2003) 2900–2918. doi:10.1016/S0006-3495(03)74712-2.

[70] Y. Shen, F. Delaglio, G. Cornilescu, A. Bax, TALOS+: A hybrid method for predicting protein backbone torsion angles from NMR chemical shifts, J. Biomol. NMR. 44 (2009) 213–223. doi:10.1007/s10858-009-9333-z.

[71] M. V. Shapovalov, R.L. Dunbrack, A smoothed backbone-dependent rotamer library for proteins derived from adaptive kernel density estimates and regressions, Structure. 19 (2011) 844–858. doi:10.1016/j.str.2011.03.019.

[72] E.F. Pettersen, T.D. Goddard, C.C. Huang, G.S. Couch, D.M. Greenblatt, E.C. Meng, T.E. Ferrin, UCSF Chimera - A visualization system for exploratory research and analysis, J. Comput. Chem. 25 (2004) 1605–1612. doi:10.1002/jcc.20084.

[73] B.R. Brooks, C.L. Brooks, A.D. Mackerell, L. Nilsson, R.J. Petrella, B. Roux, Y. Won, G. Archontis, C. Bartels, S. Boresch, A. Caflisch, L. Caves, Q. Cui, A.R. Dinner, M. Feig, S. Fischer, J. Gao, M. Hodoscek, W. Im, K. Kuczera, T. Lazaridis, J. Ma, V. Ovchinnikov, E. Paci, R.W. Pastor, C.B. Post, J.Z. Pu, M. Schaefer, B. Tidor, R.M. Venable, H.L. Woodcock, X. Wu, W. Yang, D.M. York, M. Karplus, CHARMM: The biomolecular simulation program, J. Comput. Chem. 30 (2009) 1545–1614. doi:10.1002/jcc.21287.

[74] J. Huang, A.D. MacKerell, CHARMM36 all-atom additive protein force field: Validation based on comparison to NMR data, J. Comput. Chem. 34 (2013) 2135–2145. doi:10.1002/jcc.23354.

[75] C.M. Siwy, C. Lockhart, D.K. Klimov, Is the Conformational Ensemble of Alzheimer’s Aβ10-40 Peptide Force Field Dependent?, PLoS Comput. Biol. 13 (2017) 1–26. doi:10.1371/journal.pcbi.1005314.

[76] E. Jones, T. Oliphant, P. Peterson, others, SciPy: Open source scientific tools for Python, (n.d.). http://www.scipy.org/.

[77] S. van der Walt, S.C. Colbert, G. Varoquaux, The NumPy Array: A Structure for Efficient Numerical Computation, Comput. Sci. Eng. 13 (2011) 22–30. doi:10.1109/MCSE.2011.37.

[78] S. Behnel, R. Bradshaw, C. Citro, L. Dalcin, D.S. Seljebotn, K. Smith, Cython: The Best of Both Worlds, Comput. Sci. Eng. 13 (2011) 31–39. doi:10.1109/MCSE.2010.118.

[79] J.D. Hunter, Matplotlib: A 2D graphics environment, Comput. Sci. Eng. 9 (2007) 90–95. doi:10.1109/MCSE.2007.55.

[80] W. Humphrey, A. Dalke, K. Schulten, VMD -- Visual Molecular Dynamics, J. Mol. Graph. 14 (1996) 33–38.

[81] R.J. Gowers, M. Linke, J. Barnoud, T.J.E. Reddy, M.N. Melo, S.L. Seyler, D.L. Dotson, J. Domanski, S. Buchoux, I.M. Kenney, O. Beckstein, MDAnalysis: A Python Package for the Rapid Analysis of Molecular Dynamics Simulations, in: Proc. 15th Python Sci. Conf., Scipy, 2016: pp. 102–109. http://conference.scipy.org/proceedings/scipy2016/pdfs/oliver_beckstein.pdf.

[82] N. Michaud-Agrawal, E.J. Denning, T.B. Woolf, O. Beckstein, MDAnalysis: A toolkit for the analysis of molecular dynamics simulations, J. Comput. Chem. 32 (2011) 2319–2327. doi:10.1002/jcc.21787.

[83] R.T. McGibbon, K.A. Beauchamp, M.P. Harrigan, C. Klein, J.M. Swails, C.X. Hernández, C.R. Schwantes, L.P. Wang, T.J. Lane, V.S. Pande, MDTraj: A Modern Open Library for the Analysis of Molecular Dynamics Trajectories, Biophys. J. 109 (2015) 1528–1532. doi:10.1016/j.bpj.2015.08.015.

[84] B. Han, Y. Liu, S.W. Ginzinger, D.S. Wishart, SHIFTX2: Significantly improved protein chemical shift prediction, J. Biomol. NMR. 50 (2011) 43–57. doi:10.1007/s10858-011-9478-4.

[85] A. Altis, P.H. Nguyen, R. Hegger, G. Stock, Dihedral angle principal component analysis of molecular dynamics simulations, J. Chem. Phys. 126 (2007) 1–10. doi:10.1063/1.2746330.

[86] A.P. Dempster, N.M. Laird, D.B. Rubin, Maximum Likelihood from Incomplete Data via the EM Algorithm, J. R. Stat. Soc. Ser. B. 39 (1977) 1–38. http://www.jstor.org/stable/2984875.

[87] T. Mori, N. Miyashita, W. Im, M. Feig, Y. Sugita, Molecular dynamics simulations of biological membranes and membrane proteins using enhanced conformational sampling algorithms, Biochim. Biophys. Acta - Biomembr. 1858 (2016) 1635–1651. doi:10.1016/j.bbamem.2015.12.032.

[88] N. Kucerka, M.P. Nieh, J. Katsaras, Fluid phase lipid areas and bilayer thicknesses of commonly used phosphatidylcholines as a function of temperature, Biochim. Biophys. Acta - Biomembr. 1808 (2011) 2761–2771. doi:10.1016/j.bbamem.2011.07.022.

[89] D.J. Lindberg, E. Wesén, J. Björkeroth, S. Rocha, E.K. Esbjörner, Lipid membranes catalyse the fibril formation of the amyloid-β (1–42) peptide through lipid-fibril interactions that reinforce secondary pathways, Biochim. Biophys. Acta - Biomembr. 1859 (2017) 1921–1929. doi:10.1016/j.bbamem.2017.05.012.

[90] S. Furlan, C. Hureau, P. Faller, G. La Penna, Modeling the Cu+ Binding in the 1-16 Region of the Amyloid-β Peptide Involved in Alzheimer’s Disease, J. Phys. Chem. B. 114 (2010) 15119–15133. doi:10.1021/jp102928h.

[91] S. Furlan, C. Hureau, P. Faller, G. La Penna, Modeling copper binding to the amyloid-β peptide at different pH: Toward a molecular mechanism for Cu reduction, J. Phys. Chem. B. 116 (2012) 11899–11910. doi:10.1021/jp308977s.

